# The nucleoid-associated protein IHF acts as a “domainin” protein coordinating the bacterial virulence traits with global transcription

**DOI:** 10.1101/2020.08.24.264499

**Authors:** Sylvie Reverchon, Sam Meyer, Raphaël Forquet, Florence Hommais, Georgi Muskhelishvili, William Nasser

**Affiliations:** Univ Lyon, Université Claude Bernard Lyon 1, INSA-Lyon, CNRS, UMR5240 MAP, F-69622, France; Agricultural University of Georgia, School of Natural Sciences, 0159 Tbilisi, Georgia

## Abstract

Bacterial pathogenic growth requires a swift coordination of pathogenicity functions with various kinds of environmental stresses encountered in the course of host infection. Among the factors critical for bacterial adaptation are changes of DNA topology and binding effects of nucleoid-associated proteins transducing the environmental signals to the chromosome and coordinating the global transcriptional response to stress. In this study we use the model phytopathogen *Dickeya dadantii* to analyse the organisation of transcription by the nucleoid-associated protein IHF. We determine both phenotypic effects of *ihfA* mutation on *D. dadantii* virulence and the transcriptional response under various conditions of growth. For the first time in enterobacteria, we examine the transcriptome of an IHF-depleted mutant under conditions of DNA relaxation, revealing a subtle interplay between IHF and DNA topology. We show that this mutation reorganises the genomic expression by altering the distribution of DNA supercoils along the chromosome at different length scales, thus affecting many virulence genes involved in both symptomatic and asymptomatic phases of infection, including those required for pectin catabolism. Altogether, we propose that IHF is a “domainin” protein, the inactivation of which impairs the coordination of chromosomal stress-response domains harbouring various virulence traits, thus abrogating the pathogenicity of *D. dadantii*.

## Introduction

Plant–pathogen interaction is a multifaceted process, where molecules secreted by pathogens determine both their virulence as well as success in colonizing the host. Besides having limited access to nutrients and specific oligoelements, the pathogen must cope with numerous specific challenges including exposure to various types of stress and host defence reactions. The expression of virulence and adaptive traits must therefore be tightly coordinated to ensure efficient infection.

Among the plant-pathogenic *Dickeya* species causing soft-rot disease in a wide range of hosts, one of the widely used model organisms is *D. dadantii* [1]. Studies of the invasion strategy of *D. dadantii* revealed a complex regulation combining utilization of the core metabolism and general stress-response genes with the sets of genes specifically responding to each encountered stress [2, 3]. This sophisticated genetic control mechanism is coordinated primarily at the level of transcriptional regulation of relevant genes involved in bacterial adaptation and virulence [4-8].

The bacterial adaptive response to stress involves alterations of chromosomal DNA topology modulated by abundant nucleoid-associated proteins (NAPs) that affect the expression of numerous genes, including the transcription factors [9, 10]. Changes in NAPs binding and chromosomal DNA supercoiling were shown to dynamically organise coherent domains of gene expression (aka CODOs) harbouring virulence traits, that emerge in particular constellations under conditions mimicking pathogenic growth [2, 11-13]. In *D. dadantii*, mutations inactivating either of the two highly abundant NAPs, FIS (Factor for Inversion Stimulation) or H-NS (Histone-like Nucleoid-Structuring protein), both modify the CODOs and strongly impair bacterial virulence [2, 4, 12, 14].

IHF (Integration Host Factor) is a heterodimer of IHFα and IHFβ proteins encoded by *ihfA* and *ihfB* genes, respectively, and one of the major bacterial NAPs with prominent capacity to induce sharp bends in the DNA [15]. This capacity to change the local DNA trajectory underpins the assembly of various higher-order nucleoprotein structures and facilitates long-range interactions underlying the effect of IHF not only on gene transcription, but also on site-specific DNA recombination, replication, transposition and genome packaging [16-23]. IHF was found to interact with a class of bacterial repetitive DNA elements located at the 3’ end of transcription units and was proposed to modulate gyrase binding and activity [24]. However, while the crosstalk between other NAPs and DNA supercoiling has been analysed at the whole-genome scale [11, 25], a comparable analysis is lacking for IHF.

The intracellular concentration of IHF changes with growth, increasing on transition of the cells to stationary phase [26]. In *E. coli*, IHF appears to control numerous functions including metabolic, cell cycle and adaptive processes [27, 28]. In various *Salmonella* species IHF is shown to profoundly influence the expression of virulence genes, facilitate invasion associated with transition of bacterial cells to stationary phase [29] and promote biofilm formation [30]. IHF positively regulates the expression of virulence traits in human pathogens *Vibrio cholerae* [31] and *Vibrio fluvialis* [32] as well as in plant pathogens *Erwinia amylovora* [33] and *Dickeya zeae* [34]. In the soil bacterium *Pseudomonas putida*, IHF activates the genes involved in adaptation to the post-exponential phase with limited effect on genes involved in central metabolism [35]. In spite of this eminent role in pathogenicity, the mechanistic basis for the coordinating effect of IHF on the expression of virulence and adaptation traits in bacteria remains unclear.

Previous studies revealed that in *D. dadantii* the IHF mutation leads to a growth defect and is more detrimental than in *E. coli* [36]. However, the effect of IHF on *D. dadantii* virulence has not been explored so far and in plant pathogenic bacteria in general, the contribution of IHF to global gene expression remains obscure. In this study, by using a combination of phenotypic approaches with RNA Sequencing (RNA-seq) analysis, we characterize the effect of IHF on *D. dadantii* pathogenic growth and global gene expression. We show that *ihfA* mutation modulates the CODOs, the genomic distributions of DNA supercoiling, the preferences for local organisation of transcribed units and the lagging/leading strand usage causing massive reorganization of gene expression, including virulence genes required during both the symptomatic and asymptomatic phases of infection and thus, abrogating the pathogenic growth of *D. dadantii*.

## Materials and Methods

### Bacterial strains and cell growth conditions

The bacterial strains used in this work are the *D. dadantii* strain 3937 isolated from Saintpaulia, its *ihfA* derivative [36], the *bcsA* mutant deficient in cellulose fiber production [37], the double *bcsA-ihfA* mutant obtained by generalized transduction using phage phiEC2 [38], the complemented *ihfA*/pEK strain. The pEK plasmid was generated by PCR using 3937 genomic DNA as template and the following primer couple: oligo ihf-Fw-BamHI (CCGGATCCGAATCGCCGTGATATTGCTGTGG), oligo ihf-Rev-PstI (CCCTGCAGTAATGCGGCCTTTTTGTCA). The PCR fragment was then cloned into the vector pBBR1-mcs4 between the BamHI and PstI restriction sites. In the resulting plasmid, *ihfA* is expressed under its own promoter.

All strains were grown at 30 °C either in rich Luria broth (LB) medium or in M63 minimal salts medium [39] supplemented with a carbon source (polygalacturonate (PGA) at 0.2% (w.v^-1^) and sucrose or glycerol at 0.2% (w.v^-1^)). When required, antibiotics were used as follows: ampicillin (Ap), 100 µg.mL^-1^; novobiocin (Nov) 100 µg.mL^-1^. Liquid cultures were grown in a shaking incubator (220 rpm). Media were solidified by the addition of 1.5% agar (w.v^-1^).

### Phenotype analyses

Detection of protease activity was performed on medium containing Skim Milk (12.5 g.L^-1^) and detection of cellulase activity was performed using the Congo red procedure [40]. Detection of pectinase activity was performed on medium containing PGA using the copper acetate procedure as described by [41]. Siderophore production was detected on chrome azurol S (CAS) agar plate. This assay is based on a competition for iron between the ferric complex of the dye CAS and siderophore [42]. Semi-solid agar medium containing low concentration of agar (0.4% w.v^-1^) and standard agar plates containing Carboxy-Methyl Cellulose (0.2% w.v^-1^) and glucose (0.2% w.v^-1^) were used to investigate the effect of the *ihfA* inactivation on the swimming and twitching motilities of *D. dadantii*, respectively.

For a visual assessment of cell aggregation, 5 mL of minimal M63 medium, supplemented with glycerol at 0.2% (v.v^-1^) was inoculated with an overnight culture grown in the same medium at a final density of 10^6^ cells.mL^-1^. Bacteria were grown in water bath with low shaking (55 rpm) for 48 h and cells contained in aggregates were quantified versus the total number of cells contained in both aggregate and planktonic fractions for the different strains; each value represents the mean of five different experiments and bars indicate the standard deviation.

Assay of pectate lyase was performed on toluenised cell extracts. Pectate lyase activity was determined by the degradation of PGA to unsaturated products that absorb at 235 nm [43] Specific activity is expressed as µmol of unsaturated products liberated min^-1^.mg^-1^ (dry weight) bacteria. Bacterial concentration was estimated by measuring turbidity at 600 nm, given that an optical density at 600 nm (OD_600_) of 1.0 corresponds to 10^9^ bacteria.mL^-1^ and to 0.47 mg of bacteria (dry weight) mL^-1^.

Pathogenicity assays were performed, as described in [44], with 5 × 10^6^ bacteria in 5 µl of 50 mM KH_2_PO_4_ pH 7 buffer. Assays were carried out on one hundred chicory leaves for each strain. Negative controls were performed using sterile buffer. After a 24 h incubation period, the rotted tissues were collected and weighted.

### Separation of plasmid topoisomers by gel electrophoresis

The multicopy plasmid pUC18 was extracted from *D. dadantii* WT strain and *ihfA* mutant by using the Qiaprep Spin Miniprep kit and 0.5-1 µg of plasmid DNA was electrophoresed on 1% agarose gel containing 2.5 µg.mL^-1^ chloroquine. All electrophoresis was conducted in 20 cm long agarose gel with Tris–borate EDTA (TBE) as gel running buffer. The electrophoresis was run at 2.5 V.cm^-1^ for 16 h. Under these conditions, topoisomers that are more negatively supercoiled migrate faster in the gel than more relaxed topoisomers. Distribution of topoisomers was analysed using Image Lab 6.0 software (Biorad).

### RNA extraction and transcriptomics data

The wild type (WT) strain and its *ihfA* derivative were used to analyse the global gene expression of cells grown under different conditions: M63 supplemented with 0.2% (w.v^-1^) sucrose as carbon source, with or without 0.2% (w.v^-1^) PGA. Cells were grown to the early exponential phase (OD_600_ = 0.2) and to the early stationary phase (OD_600_ = 0.9-1.2 for cells grown in M63/sucrose, and OD_600_ = 1.9-2.2 for cells grown in M63/sucrose/PGA medium). The different OD_600_ retained for the various culture media and strains correspond to a similar growth stage (*i*.*e*., transition from late exponential phase to early stationary phase), then aliquots were transferred to two flasks. One of them was kept as a control and the second was treated for 15 min with novobiocin to 100 µg.L^-1^ final concentration. At this concentration, novobiocin has no impact on *D. dadantii* parental strain growth [14]. For each condition, total RNAs were extracted using the frozen-phenol procedure [45]. Control experiments for RNA extraction quality, absence of DNA contamination, and qRT-PCR validation of selected genes were conducted as previously [11]. Further steps were carried out by Vertis Biotechnologie AG (http://www.vertis-biotech.com): rRNA depletion using the Illumina Ribo-Zero kit, RNA fragmentation, strand-specific cDNA library preparation, and Illumina NextSeq500 paired-end sequencing (∼15 million paired reads per sample). Transcriptomes were analysed as previously described [46] using softwares FastQC, Bowtie2 (reference genome NC 014500.1), and DESeq2 for differential expression. Significantly activated/repressed genes were selected with a threshold of 0.05 on the adjusted p-value.

### Computational and statistical methods

The predicted binding sites of IHF were generated from its position weight matrix with Prodoric2 [47]. Orientational distributions were analysed with a homemade Python code, where a gene/region is defined as convergent/divergent/tandem based on the orientation of the two neighbouring genes. Proportions were compared with chi-square tests. For genome-wide analyses, all parameters were computed over 500 kb scanning windows shifted by 4 kb, resulting in 1230 overlapping windows. The numbers of activated/repressed genes and of predicted binding sites in each window were transformed into z-scores by comparison to the null hypothesis of a homogeneous distribution of the considered quantity over the chromosome. Melting energies were computed with the parameters of Santa Lucia [48] and averaged over each window.

## Results

While the *ihfB* mutation severely compromises the growth of *D. dadantii* 3937 [36], we took advantage of the *ihfA* mutation to disable the formation of the IHF heterodimer in *D. dadantii* cells. In *E. coli*, it was observed that, potentially, both IHFα and IHFβ could form homodimers capable of binding DNA and supporting the lambda phage site-specific recombination reaction *in vitro* [49]. However, the binding affinity of the IHFβ homodimer is by two orders of magnitude lower than that of the IHF heterodimer, whereas IHFα homodimer is highly unstable [50, 51]. In *D. dadantii*, we observed that the *ihfA* mutant is able to grow and achieve, albeit with delay, reasonably high cell densities (Fig 1A), making them amenable to experimental investigation. Given the fact that the lack of IHF is more detrimental for *D. dadantii* than for *E. coli* [36] and that the *ihfB* gene expression is unaffected in *ihfA* mutant (Table S2 in Supp. Mat.), as also observed in *E. coli* [33], it is conceivable that bacterial growth is permitted by the activity of the low-affinity IHFβ homodimer. It is also noteworthy that in *Salmonella enterica*, the single *ihfA* and *ihfB* mutants and the double *ihfA/ihfB* mutant are all able to grow, and exhibit similar transcriptional responses [29]. In this study, we assume that all alterations induced by *ihfA* mutation are due to the lack of the IHF heterodimer in the cell.

**Fig. 1:**
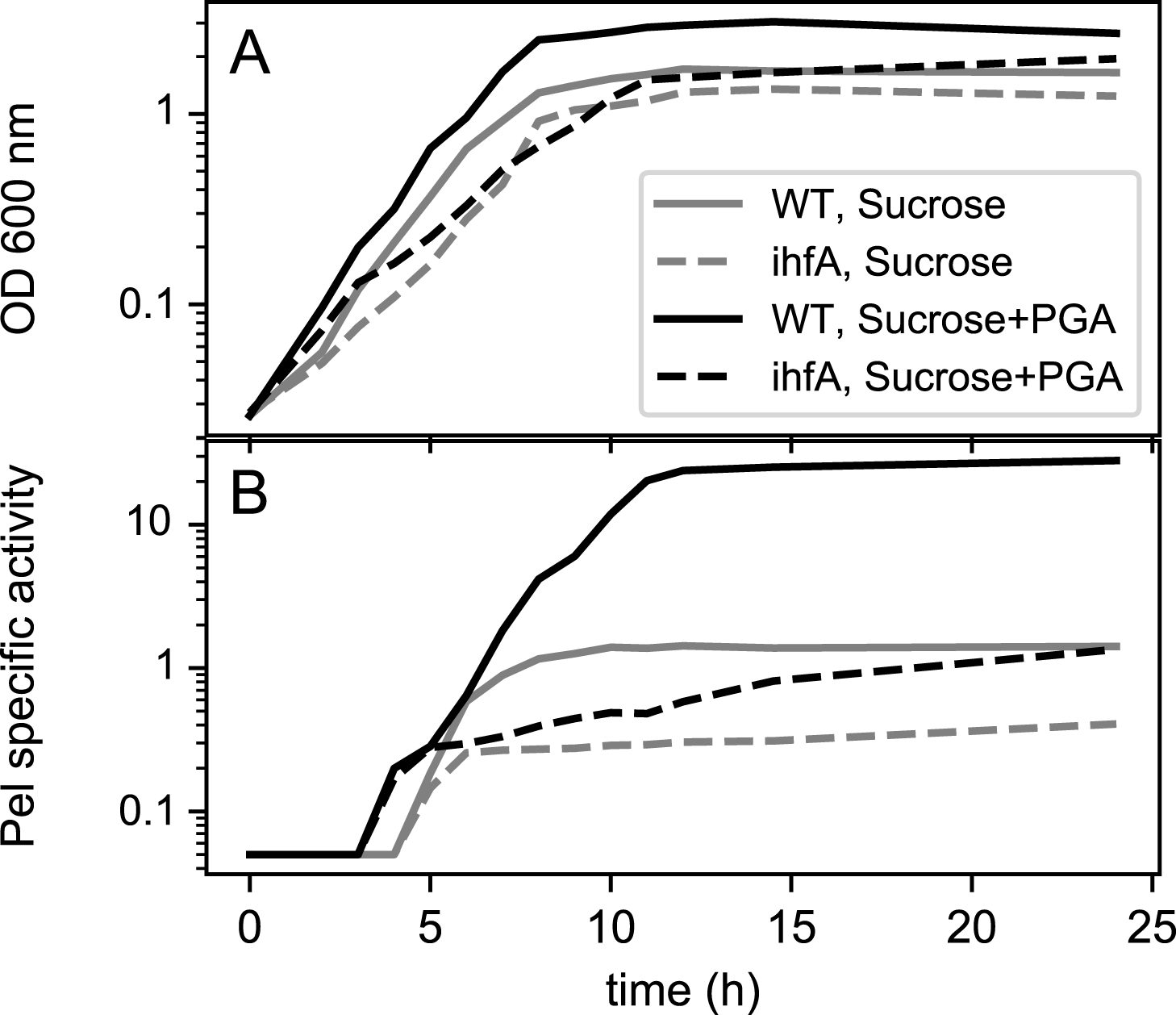
(A) Growth and (B) pectate lyase production of *Dickeya dadantii* 3937 wild type strain (solid lines) and its *ihfA* derivative (dashed lines) grown in liquid M63 minimal medium supplemented with sucrose (gray) or sucrose+polygalacturonate (black). Samples were taken every hour. Pel specific activity is expressed as µmol of unsaturated product liberated per min per mg of bacterial dry weight. The experiment was repeated three times and curves from a representative experiment are shown (variation between experiments was less than 15%).

### IHF regulates cell motility, production of plant cell wall degrading enzymes, siderophores and cell aggregation

Inactivation of *ihfA* resulted in reduced production of many virulence factors such as pectinases (mainly pectate lyases), cellulases and proteases responsible for the destruction of the plant cell wall and production of the soft rot symptoms (Fig. 2 ABC). The *ihfA* mutation also reduced the production of siderophores (Fig. 2D) required for the systemic progression of maceration symptoms in the hosts [52, 53]. Furthermore, the swimming and twitching capacities of *D. dadantii* essential for searching favourable sites of entry into the plant apoplast were reduced (Fig. 2E, F). Complementation of the *ihfA* mutant with the *ihfA* gene carried on the pEK plasmid restored the WT phenotype in all cases, indicating that inactivation of *ihfA* was responsible for impaired functions (Fig 2).

**Fig. 2:**
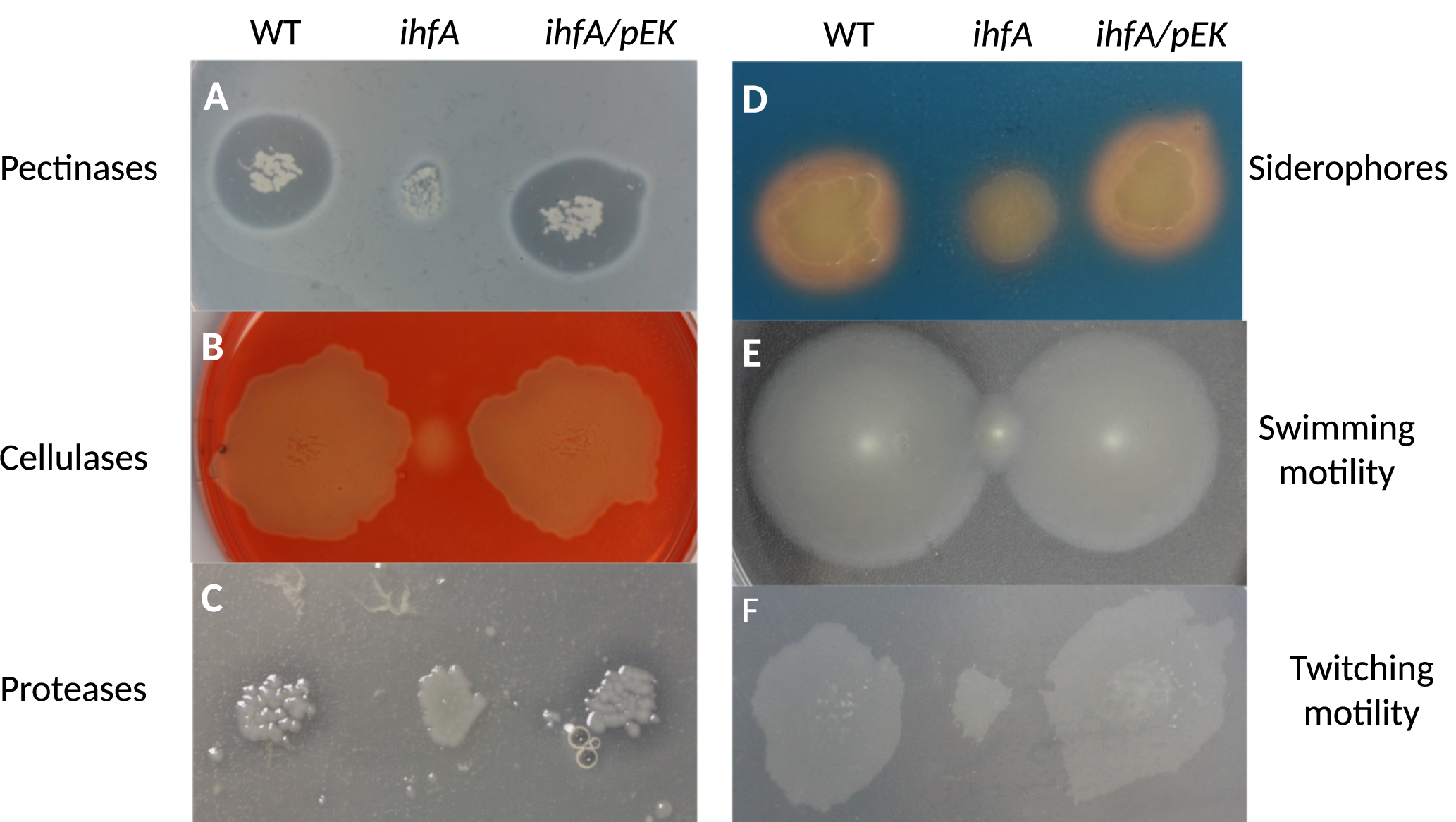
Phenotypes of the *D. dadantii ihfA* mutant and its complemented derivative *ihfA*/pEK. **(A)** Pectinase activity was observed after growth on PGA-containing plate and staining with copper (II) acetate solution. **(B)** Cellulase activity was detected after growth on Carboxy-Methyl Cellulose-containing plate, followed by staining with Congo red. **(C)** Protease activity was revealed, after growth on skim milk-containing plate by a translucid halo surrounding the bacteria growth area. **(D)** Siderophore production was detected after growth on chrome azurol S (CAS) agar plate. **(E)** Swimming motility inside the soft agar medium, was estimated by picking bacterial colonies with a thin rod inside a semi-solid agar plate containing low concentration of agar (0.4% w.v^-1^). **(F)** Twitching motility at the surface of the agar medium was estimated by picking bacterial colonies with a thin rod inside a standard agar plates containing Carboxy-Methyl Cellulose (0.2% w.v^-1^) and glucose (0.2% w.v^-1^). Plates were incubated at 30°C for 24 h before measuring colony expansion.

*D. dadantii* produce cellulose fibrils, which enable the cells to aggregate on the plant surface and form a biofilm resistant to desiccation [37]. The cellulose fibrils are not required anymore when bacteria enter into the plant apoplast and colonize this inter-cellular compartment by using their motility function. Cell aggregate formation was analysed in the WT and *ihfA* strains grown under low shaking conditions (Fig. 3). We found that the growth medium of both the WT strain and the complemented *ihfA* mutant appeared turbid and at the bottom of the tubes, only small cell aggregates were observed (Fig. 3A). In contrast, the growth medium of the *ihfA* mutant was clear and a large cell aggregate was observed (Fig. 3A). The estimated percentages of cells in aggregates were 29% for the WT strain, 37% for the complemented *ihfA* strain and 76% for the *ihfA* mutant (Fig. 3B). Inactivation of the *bcsA* gene encoding cellulose synthetase suppressed the aggregation phenotype observed in the *ihfA* mutant. Taken together, these results demonstrate that IHF negatively regulates the production of cellulose fibril adherence structures.

**Fig. 3:**
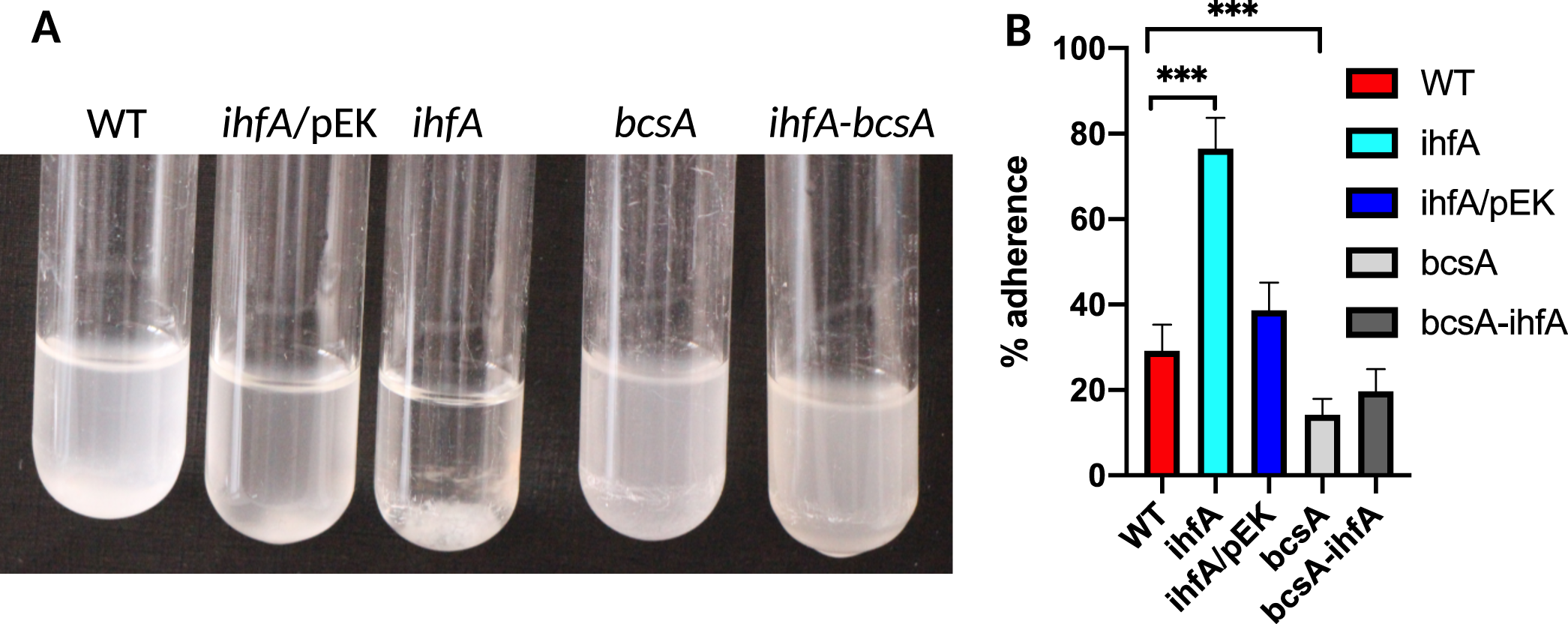
**A**. Aggregation phenotype of the *D. dadantii* 3937 WT strain and its derivatives, *ihfA, bcsA, ihfA-bcsA* and the complemented *ihfA*/pEK strain. Strains were grown in minimal M63 medium supplemented with glycerol for 48 h under low shaking condition (55 rpm). **B**. Quantification of the cells present in the aggregates and in the planktonic fractions for the different strains; each value represents the mean of five experiments and bars indicate the standard deviation. *** indicates a significant difference relative to the WT (*P* < 0.001, Mann-Whitney test).

### Inactivation of *ihfA* strongly reduces pectinase activity and impairs *D. dadantii* virulence

Since the soft rot symptoms of infection by *D. dadantii* are mainly due to the production and secretion of pectate lyases (Pels), we monitored the production of Pels and their induction by polygalacturonic acid (PGA, a pectin derivative) in both *ihfA* and WT cells.

As mentioned above, compared to the WT strain, the *ihfA* mutant exhibited a slower growth rate, reaching lower maximal densities both in presence and absence of PGA (Fig. 1A). In the *ihfA* mutant, the production of Pels was strongly reduced at all sampling times both in PGA-induced and non-induced growth condition, while the growth phase-dependence of Pel production was retained (Fig. 1B). However, the most striking effect was the strongly reduced induction by PGA, as the production of Pels was weaker in the *ihfA* mutant in presence of PGA than in the WT strain in absence of inducer (but it kept increasing slowly due to the low remaining Pel enzymatic activity). Since the induction by PGA itself is dependent on Pel production via a positive feedback loop [7] these results suggest that *ihfA* mutation affects the expression of *pel* genes by two separate mechanisms: (1) inducer-independent transcriptional repression by a factor of ∼5 (as observed in sucrose medium), and (2) decrease of *pel* sensitivity to PGA induction, incurring an additional factor of ∼5 (Fig. 1B)

Altogether, these observations suggest that IHF plays a central role in *D. dadantii* pathogenicity by activating several key virulence factors. We therefore analysed the impact of *ihfA* mutation on *D. dadantii* virulence, using the chicory leaf assay (Fig. 4). As expected, the pathogenic growth is drastically reduced when the plant is infected with the mutant strain. Complementation of the *ihfA* mutation with the *ihfA* gene carried on the pEK plasmid fully restored the virulence. This result indicates that IHF is required for the development of soft rot disease induced by *D. dadantii* in chicory.

**Fig. 4:**
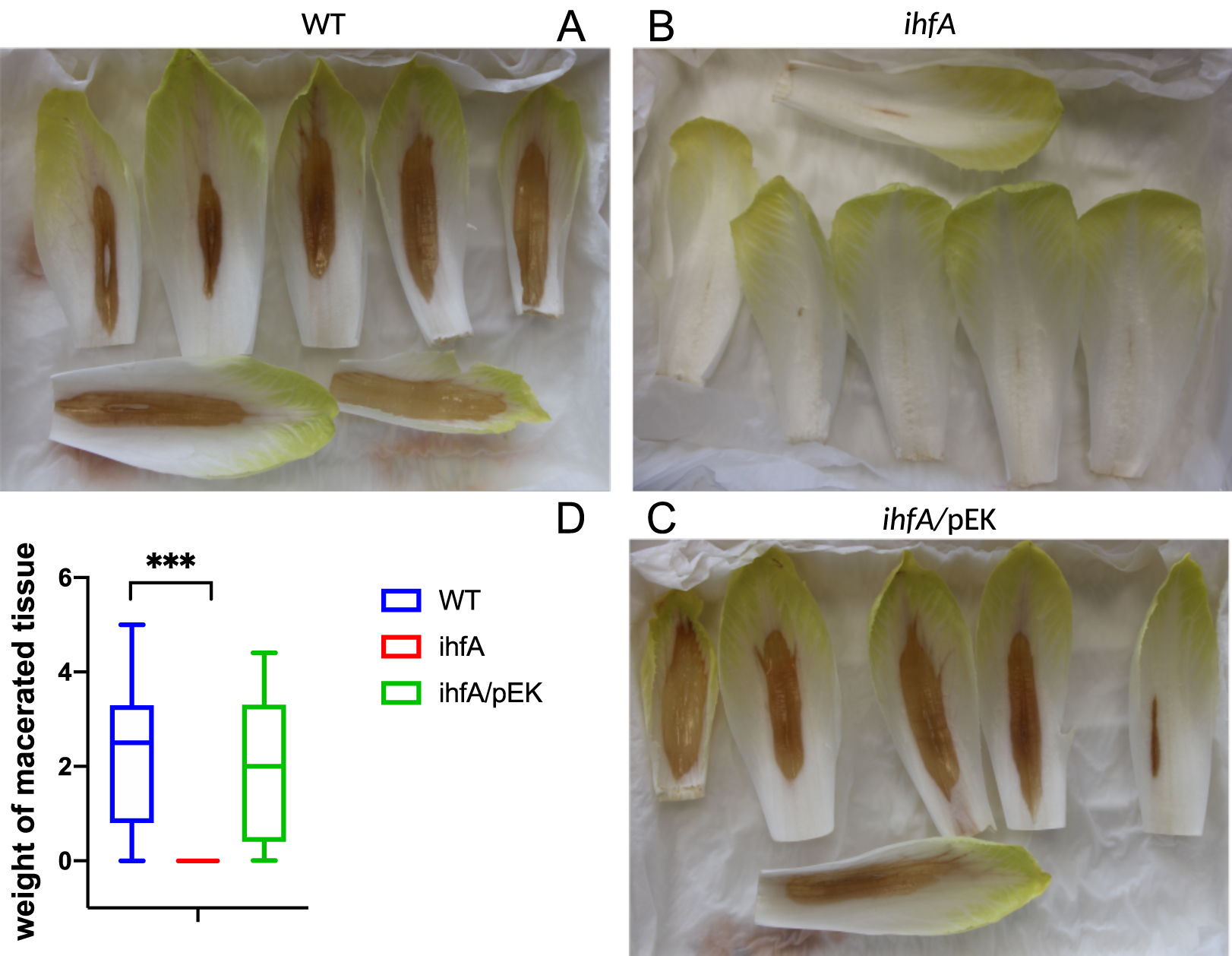
Maceration capacity of WT, *ihfA* and *ihfA* complemented (*ihfA*/pEK) strains on chicory leaves. Representative specimen of chicory leaves infected by the wild-type strain **(A)**, the *ihfA* mutant **(B)**, the complemented *ihfA*/pEK strain **(C)**. One hundred chicory leaves were infected for each strain, using 5 µl of bacterial suspension (10^8^ cfu.mL^-1^ in 50 mM KH_2_PO_4_ pH 7 buffer). After incubation in a humid chamber for 24 h at 30°C, the weight of macerated tissue was measured. **(D)** Boxplot from 100 data points. The calculated median value is 2.5 g of macerated tissue for the WT strain, 0 g for the *ihfA* mutant and 1.9 g for the complemented *ihfA*/pEK strain. Centre lines show the medians; box limits indicate the 25th and 75th percentiles. *** indicates a significant difference relative to the WT (*P* < 0.001, Mann-Whitney test).

### The *ihfA* mutation reorganizes the *D. dadantii* transcriptome

As seen above, the strong phenotypic effect of the *ihfA* mutation results from the inactivation of several key virulence functions. To broaden this observation and assess the role of IHF in global gene regulation, we analysed the *D. dadantii* transcriptome by RNA-seq under various growth conditions: in early exponential phase in M63 minimal medium supplemented with sucrose as carbon source, and at transition to stationary phase in M63 supplemented with either sucrose or sucrose + PGA (see Materials and Methods).

The most dramatic effect of IHF on gene regulation was observed when the transcriptomes were compared in presence vs absence of PGA, in both *ihfA* and wild type cells. Whereas 762 and 752 genes were found respectively up-and down-regulated by PGA in the wild-type strain, only about a dozen of genes were altered in the *ihfA* background under the same conditions (Supplementary Table S1). This finding is fully consistent with the impaired cell wall-degrading enzyme activity in *ihfA* strain described above (Fig. 1B and Fig. 2) and indicates that the mutant almost completely lost the ability to metabolise this important energy source in the medium.

In order to analyse the regulatory effect of IHF, we next focused on the transcriptomes obtained with sucrose as sole carbon source (Supplementary Table S2). We found that 809 genes in total were differentially expressed during the exponential and stationary phases in the WT vs *ihfA* transcriptome comparisons (Fig. 5A). This large number exceeds that of the differentially expressed genes reported in *E. coli* using DNA microarray analysis [28], and is consistent with the notion that IHF acts as a global regulator in *D. dadantii* 3937. Among these genes, 432 were differentially expressed during exponential growth and 682 during stationary phase, consistent with the major impact of IHF at this stage of growth [26]. Out of the 809 differentially expressed genes, 365 fell into the category of “IHF-repressed” (*i*.*e*. up-regulated in *ihfA* mutant) and 444 fell into the category of “IHF-activated” (*i*.*e*. down-regulated in *ihfA* mutant).

**Fig. 5:**
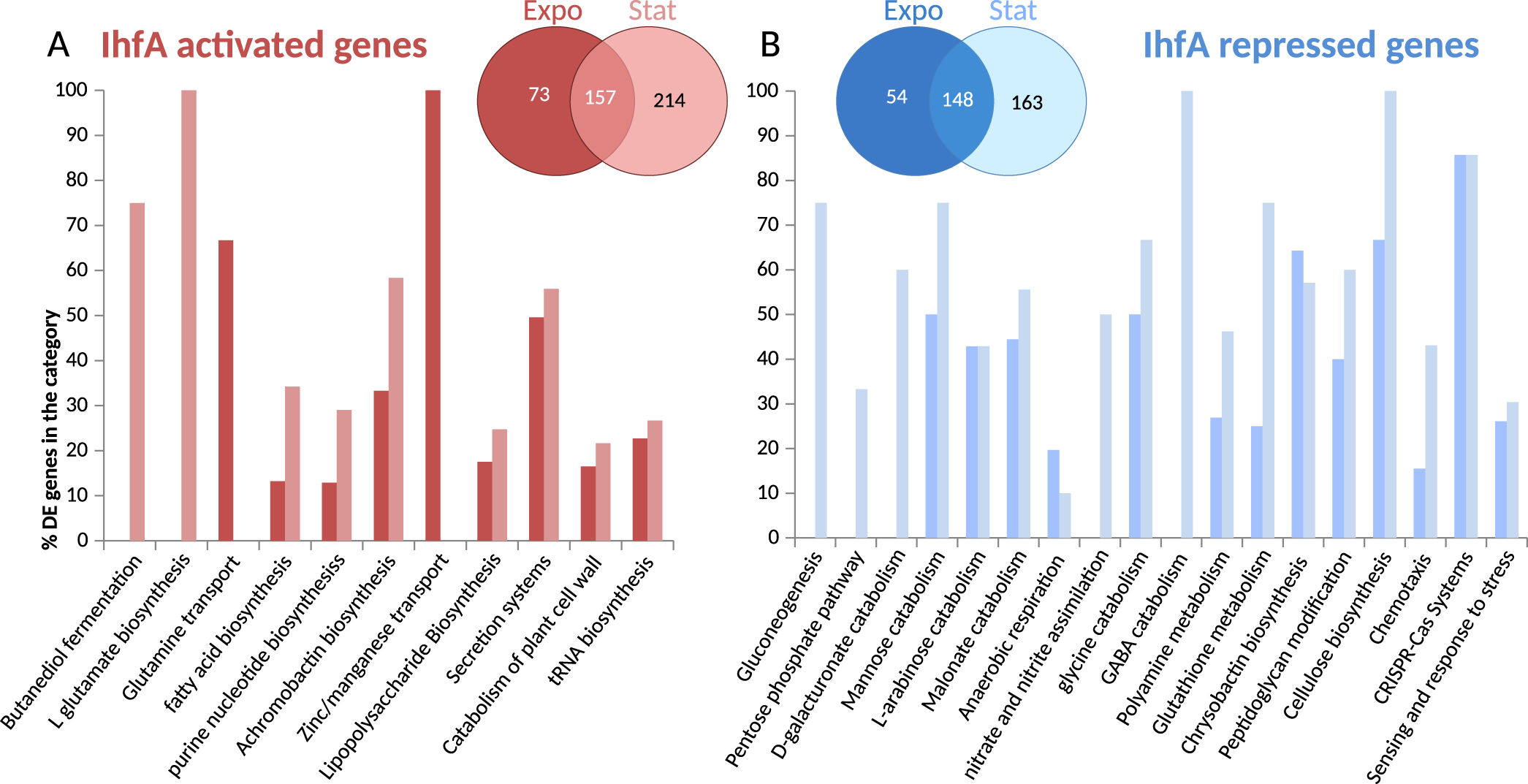
Transcriptional response to *ihfA* mutation. Venn diagram and functional repartition of genes significantly activated (A) or repressed (B) in the *ihfA* vs WT strain. Only genes with an adjusted p-value < 0.05 and fold-change > 2 (either positive or negative) were considered.

### Functional classes of differentially expressed genes

An analysis of the biological functions significantly enriched among differentially expressed genes in the *ihfA* mutant (Fig. 5) suggests a global reorganization of the central metabolism, including carbon storage via gluconeogenesis, use of alternative carbohydrate (galacturonate, arabinose, mannose) catabolic pathways and reduction in the biosynthesis of fatty acids and nucleotides. Such observations are indicative of a profound modification in carbon flow. A preference towards anaerobic metabolism was observed, with an increase in malonate catabolism, increase in the pentose phosphate pathway in connection with the capacity to regenerate the redox potential (NADPH) of cells. Nitrogen metabolism was also affected with an increased assimilation of nitrate and nitrite as well as an increase in the catabolism of amino acids (Glycine and GABA catabolism) and polyamines. This profile is consistent with the role of IHF in regulation of RpoN-dependent genes [18, 54] and nitrogen regulation, as also observed in *Klebsiella pneumoniae* [55] and *Rhizobium meliloti* [56].

In *ihfA* mutant cells, numerous virulence functions were down-regulated, including the secretion systems Prt T1SS, Out T2SS, Hrp T3SS, T6SS as well as their effectors. Genes encoding plant cell wall-degrading enzymes including proteases PrtABCG secreted by T1SS, pectinases Pels, cellulase Cel5Z, xylanase XynA secreted by T2SS were down-regulated as well as other type 2 effectors such as the extracellular necrosis inducing protein NipE and the two proteins AvrL-AvrM favouring disease progression. Genes encoding the type 3 effectors (DspE, HrpN, HrpW) which suppress plant immunity and promote pathogenesis were down-regulated as well as the genes encoding type 6 effectors (RhsA, RhsB) which have been identified as antibacterial effectors but may have key functions within the plant host [57].Furthermore, the *rhlA* gene responsible for biosurfactant biosynthesis promoting plant surface colonization was down-regulated [6]. Biosynthesis of the two siderophores, which allow bacteria to cope with the restricted iron bioavailability in the plant [52, 58, 59] and which also manipulate plant immunity [53] was distinctly affected in *ihfA* mutant: achromobactin, which is produced when iron becomes limiting was down-regulated, whereas chrysobactin prevailing under severe iron deficiency was up-regulated. In addition, *ibpS* and *indABC* genes responsible for production of an iron scavenging protein [60] and of the antioxidant pigment indigoidine [61] respectively, were down-regulated. Regarding motility, the *pil* genes involved in twitching motility were down-regulated, but the flagellar genes were not significantly affected. Also the genes encoding several important regulators of virulence were affected: the repressor gene *pecT* was up-regulated while the nucleoid-associated protein gene *fis* was down-regulated, as well as the genes of regulators *hrpL, mfbR, fliZ* and the small non-coding RNA *rsmB* (Supplementary Table S2).

Overall, the decrease in anabolic functions concomitant with an increase in catabolic functions provides a signature of the “survival mode” behavior of the *ihfA* mutant, consistent with modifications of the bacterial envelope (peptidoglycan, outer-membrane porins, lipopolysaccharide, exopolysaccharide in particular cellulose), and with an increased stress response and activation of the CRISPR-Cas defences.

We shall note however, that the number of genes among these enriched functional groups represented only a half of the total number of differentially expressed genes. The other half was more dispersed among various pathways, making it difficult to allocate them to particular functions.

### IHF binding predictions at gene promoters

We next attempted to get an insight into the mechanism underlying the regulatory effect of IHF. The latter is known to act as a global transcription factor binding at many gene promoters but, in contrast to the closely related NAP HU, IHF recognizes a relatively well-defined sequence motif (consensus WWW**TCA**ANNNN**TT**R), possibly related to the extreme bend of about 180° that it introduces in the DNA [15]. We took advantage of this feature to predict the distribution of potential IHF binding sites in the chromosome, and determine which responsive genes are possibly regulated through a direct activation or repression of their promoter by IHF. The results are provided in Table S5 in Supplementary data, with predictions filtered with a loose threshold (5000 putative binding sites retained). Around 36% of the 809 differentially expressed genes have an upstream IHF binding signal, with a weak but significant enrichment compared to the proportion obtained with random sets of genes of identical size (31% in average, p-value=0.007). Keeping only 500 sites with strongest scores reduced this proportion to 8%, but the relative enrichment became stronger (5% for random genes, p-value=0.003).

This limited overlap between the identified IHF regulon and the set of genes with binding signals may partly reflect the limitations of the prediction method, but is not unexpected based on the properties of the protein. On the one hand, in contrast to most transcriptional regulators targeting a small number of promoters, IHF is a “hub” of the regulatory network and thus activates or represses many other genes in an indirect manner. On the other hand, IHF is known to act not only as a “digital” transcription factor, but also as a NAP that binds DNA with loose specificity, and modulates the transcriptional activity by an “analogue” mechanism [62] involving global modifications of the chromosome architecture. It is the latter mechanism that we now investigate in more detail.

### Regulatory interplay between IHF and DNA supercoiling

One of the major factors in bacterial chromosome compaction and analogue transcriptional regulation is DNA supercoiling, which is controlled by a crosstalk between topoisomerase enzymes and nucleoid-associated proteins [10]. We therefore addressed the question of a coupling between the regulatory action of IHF and changes of DNA topology. Such an effect was shown to underpin the IHF regulation of the *ilvP* promoter of *E. coli* [63], but was never investigated at genomic scale in an enterobacterium.

We assessed the effect of IHF on DNA topology by high-resolution agarose gel-electrophoresis of pUC18 plasmids isolated at different stages of growth from both WT and *ihfA* mutant strains grown with or without treatment by the gyrase inhibitor novobiocin (Fig. 6). Whereas addition of novobiocin induced relaxation of DNA in both strains, the *ihfA* mutant did not exhibit any significant change in DNA topology compared to the WT strain. This observation is consistent with the lack of variation in topoisomerase gene expression in the *ihfA* transcriptome (Table S2) and can be related to previously observed similar contrasting effects of *fis* and *hns* mutations in *D. dadantii* and *E. coli*[14] [25].

**Fig. 6:**
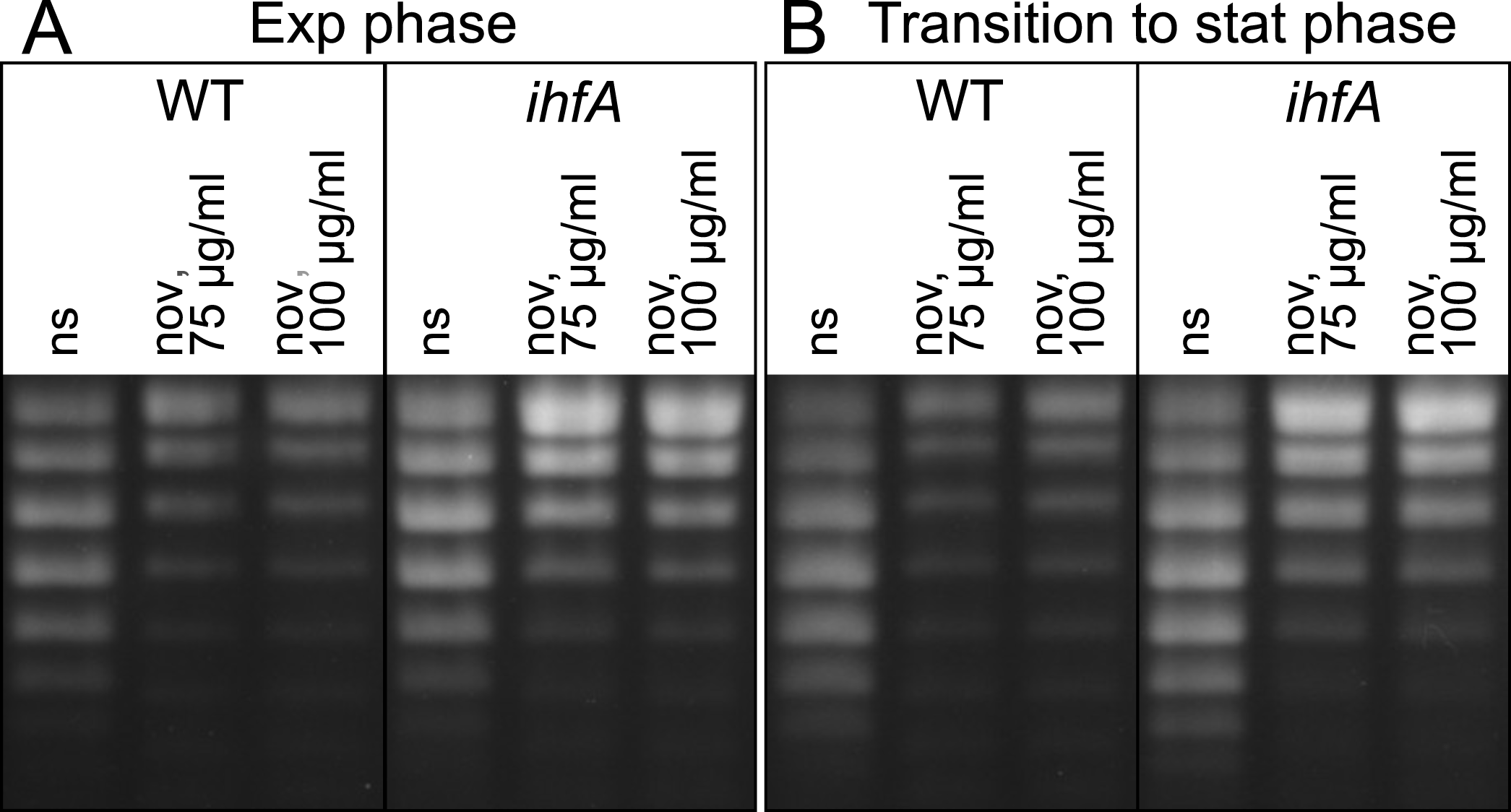
DNA supercoiling in the *D. dadantii* WT strain and its *ihfA* derivative. Topoisomers of plasmid pUC18 were isolated and separated on agarose gel containing 2.5 µg.mL^-1^ chloroquine. At this concentration, the more negatively supercoiled topoisomers migrate faster in the gel. The experiment was performed three times and a typical result is shown. Growth phases and novobiocin treatment are indicated (ns, none stressed cells).

Given that *ihfA* mutation has no noticeable impact on global topology of plasmid DNA, is it equally irrelevant with regard to the supercoiling response of chromosomal genes? To answer this question, we monitored the transcriptional response of the WT and mutant strains to novobiocin shock. The number of genes significantly activated or repressed in response to novobiocin addition was slightly lower in *ihfA* mutant (774 in exponential phase, 809 on transition to stationary phase) than in the WT strain (914 and 1133 genes, respectively). Strikingly, however, the genes sensitive to gyrase inhibition in the two strains were mostly different (Fig. 7A). Furthermore, the effect of the *ihfA* mutation differed depending on the supercoiling level of the DNA, with many genes responding specifically in the relaxed state of the chromosome (Table S4 in Supplementary Data). A set of functional pathways belonging to sugar catabolism (ribose, lactose), amino-acid biosynthesis (leucine, valine, isoleucine, tryptophan, histidine), malate and ureide catabolism, flagellar assembly and disulfide bond formation was enriched in this group. Among the plant cell wall-degrading enzymes, the galactan catabolism was specifically repressed (Fig. 7B).

**Fig. 7:**
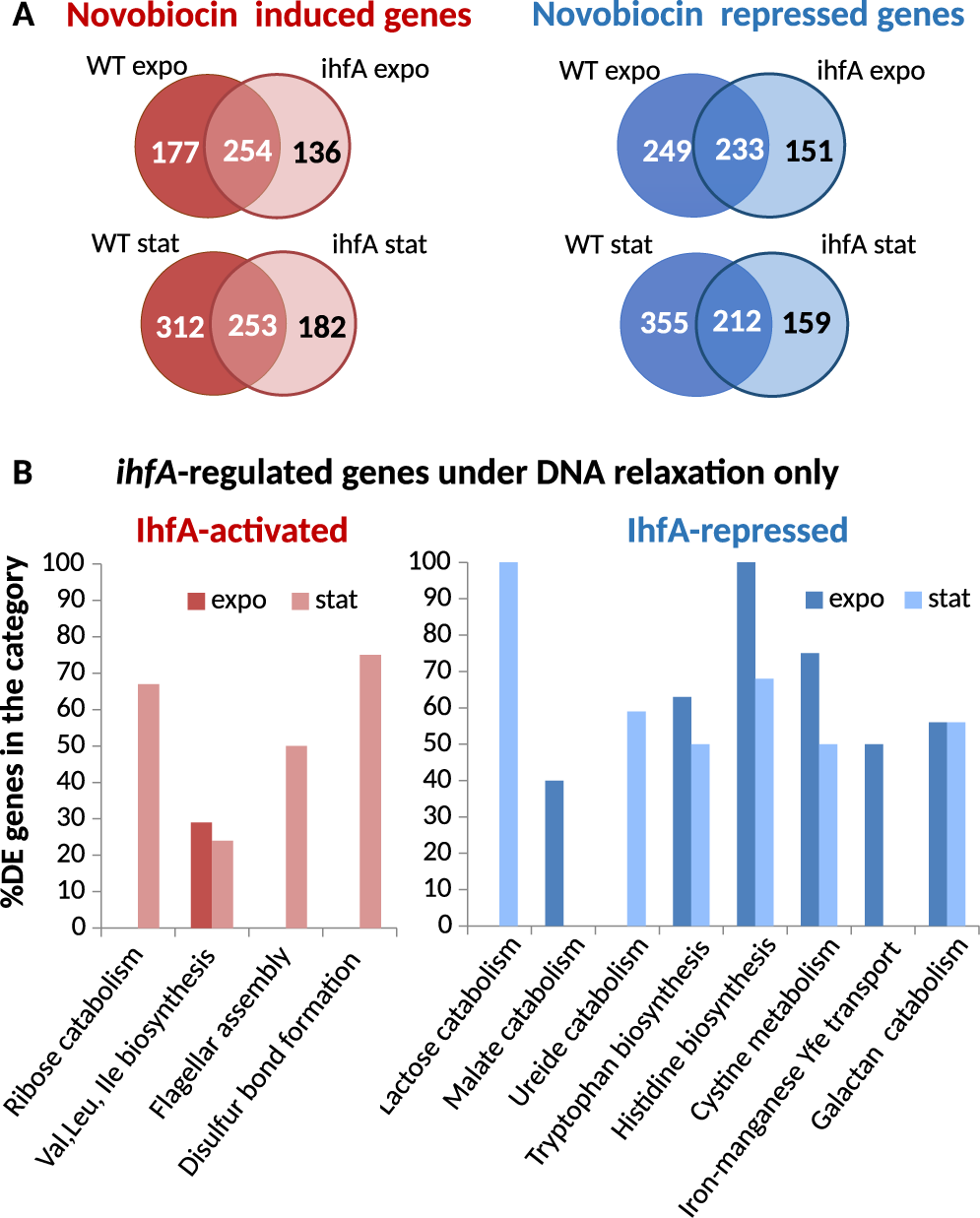
Transcriptional response to global DNA relaxation induced by novobiocin treatment of the *D. dadantii* WT and *ihfA* strains. **A**. Venn diagrams of significantly activated and repressed genes after a novobiocin shock, either in the WT or *ihfA* strains, in exponential and stationary phases. Only the significant differentially expressed genes (p-value<0.05, Fold-Change>2 or Fold-Change < 0.5) were considered. **B**. Functional gene groups significantly enriched among the *ihfA*-regulated genes under conditions of DNA relaxation. A statistical enrichment analysis was carried to extract biological processes significantly over-represented in the set of *ihfA*-regulated genes under conditions of DNA relaxation.

This nontrivial response to DNA relaxation by novobiocin indicates that, although the *ihfA* mutation apparently does not change the *global* supercoiling level of DNA, it has a strong impact on how gyrase affects transcription at the genomic scale, which, in turn, likely reflects a profound modification of the *local* distribution of DNA supercoiling along the chromosome. We now analyse this phenomenon at two successive length-scales.

### Orientational organisation of IHF transcriptional regulation by genomic architecture

Gyrase inhibition is known to affect the gene expression globally, but with a bias for convergent genes, *i*.*e*., genes located on complementary DNA strands and facing each other [46]. This effect is related to the asymmetric build-up of positive and negative supercoils during the transcription process itself, which underpins an intimate regulatory coupling between transcription and DNA supercoiling and affects neighbouring genes differently depending on their relative orientation [46]. Interestingly, whereas DNA relaxation by novobiocin in WT cells reduces the utilisation of divergent genes, the lack of IHF heterodimer completely reverses this effect, with divergent genes being more activated than convergent ones in the *ihfA* mutant (Fig. 8A). This means that IHF plays a crucial role in organising the DNA supercoils in the vicinity of transcribed genes. The relation between gene orientation and IHF regulation can be also analysed by comparing the gene expression of the two strains (Fig. 8B). Whereas the presence of IHF already favours convergent genes in standard growth conditions, the effect is considerably enhanced in a novobiocin-relaxed chromosome, with more than a two-fold higher proportion compared to divergent genes. This effect likely results from the combination of two factors: (1) divergent regions are more AT-rich, and thus tend to attract IHF more favourably than convergent ones, as visible in the distribution of predicted binding sites (Fig. S1 in Supplementary Materials); (2) IHF is known to recognise the structural properties (geometry) of DNA as well as its sequence [64] and its binding might thus also be modulated differently by DNA supercoils resulting from adjacent transcription in these regions. This differential effect of IHF on the convergent and divergent transcription units raised the question about its possible influence on the directionality of genomic transcription.

**Fig. 8:**
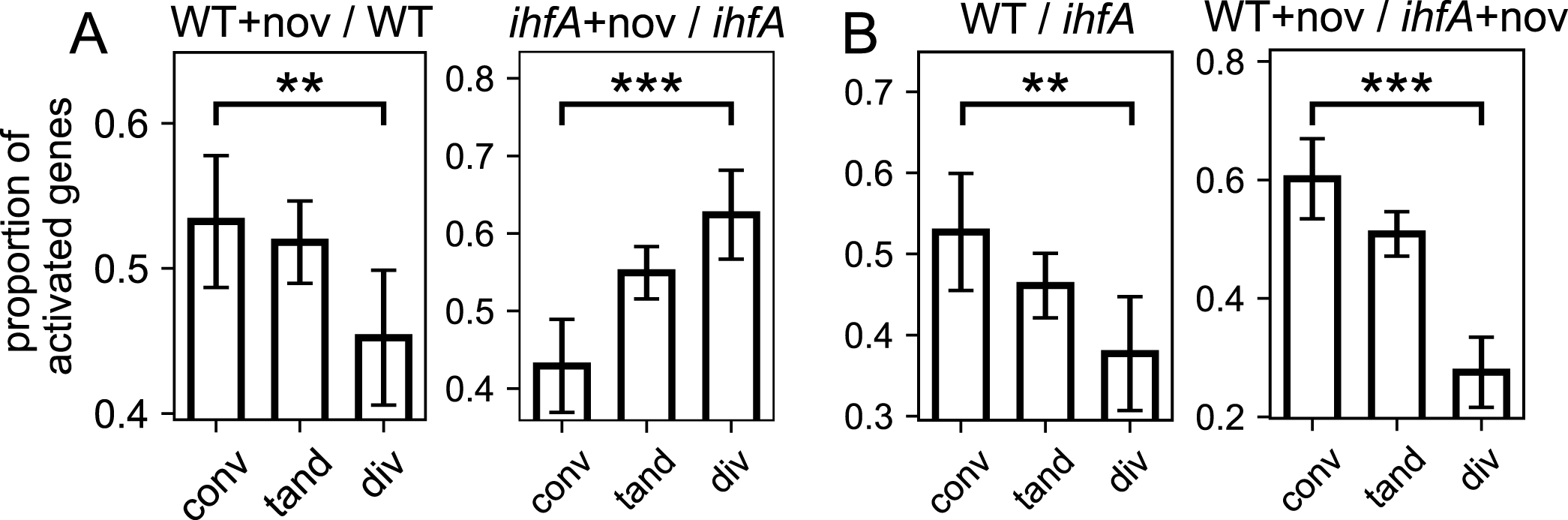
Local organisation of IHF transcriptional regulation by genomic architecture. (A) Proportion of activated genes among convergent, tandem and divergent differentially expressed genes during gyrase inhibition by novobiocin, in the WT strain and its *ihfA* derivative. The selective activation of convergent vs divergent genes is completely reversed in the absence of IHF. (B) Comparison of the transcriptional effect of *ihfA* mutation to WT depending on gene orientation, in absence and presence of novobiocin. All proportions are computed at the transition to stationary phase. Error bars represent 95% confidence intervals.

We therefore analysed the preferences of leading and lagging strand utilisation in the WT and *ihfA* mutant cells. We found that DNA relaxation by novobiocin addition has no effect on leading versus lagging strand utilization, whether in the WT or in the *ihA* mutant cells (Fig. 9A), showing that gyrase is not specifically required for transcription along one of them. On the other hand, when we compared the expression of genes between the WT and *ihfA* mutant cells, we found that on average, the leading strand utilization was significantly preferred in the WT cells, and this effect was enhanced under conditions of DNA relaxation (Fig. 9B). Thus, IHF imposes directionality on genomic transcription and facilitates leading strand utilization. This suggests that *ihfA* mutation may cause a global reorganization of genomic transcription in response to changes of DNA supercoiling and especially, affect the organization of CODOs.

**Fig. 9:**
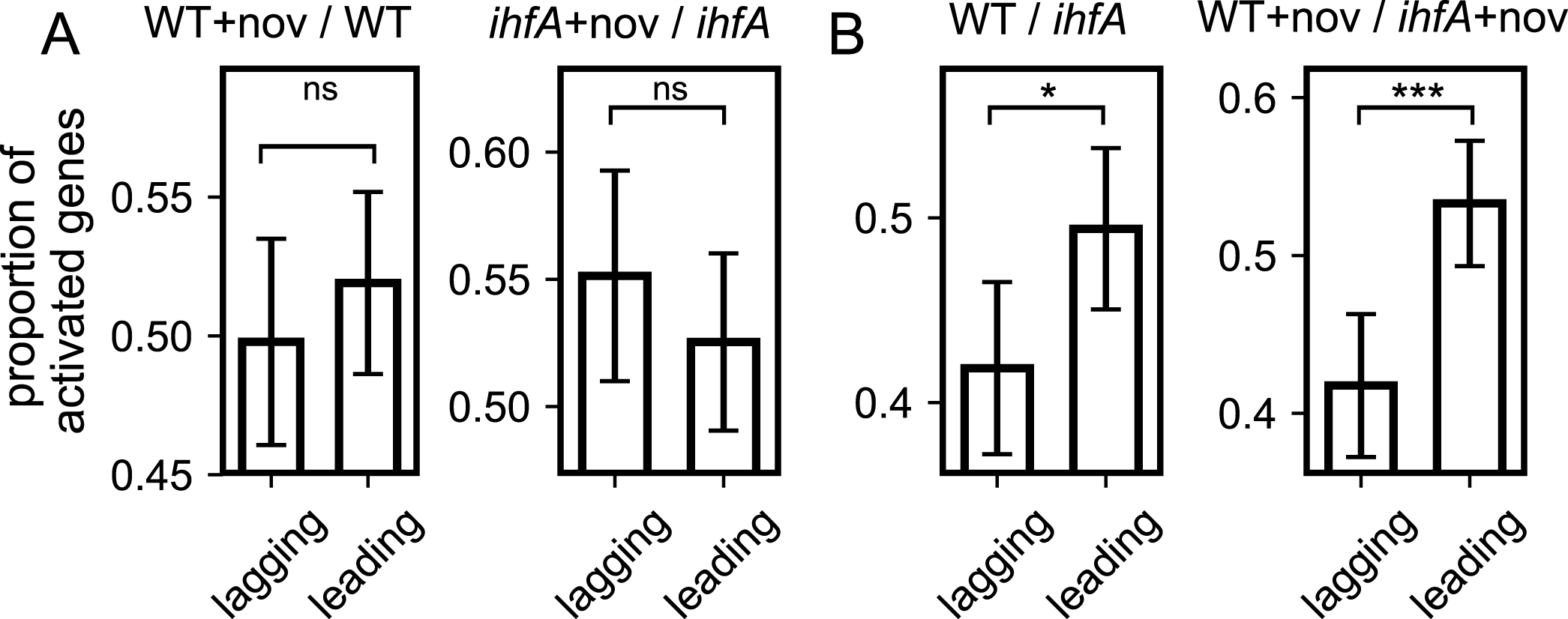
Effect of IHF on the leading versus lagging strand utilisation. (**A**) Proportion of activated genes among the differentially expressed genes on the leading and lagging strand during gyrase inhibition by novobiocin, in the WT strain and its *ihfA* derivative. Note the selective activation of lagging strand transcription in the absence of IHF. **(B)** Comparison of the transcriptional effect of *ihfA* mutation to WT on the genes expressed on the leading and lagging strand in absence and presence of novobiocin. All proportions are computed at the transition to stationary phase. Error bars represent 95% confidence intervals.

### Global organisation of IHF transcriptional regulation in chromosomal domains

We now enlarge the scanning window of our analysis, in order to describe how IHF regulation is distributed along the chromosome (Fig. 10). At that scale, the genome of *D. dadantii* exhibits a symmetrical organisation into four large regions [11] of various G/C contents or DNA thermodynamic stabilities (denoted SD1, SD2, LD1, LD2, outer wheel of Fig. 10A), which are highly conserved among *Dickeya* species [13]. The predicted binding of IHF (second outer wheel) follows the same pattern, which reflects the protein’s preference for AT-rich sequences but may also entail a relation between DNA physical properties and the binding and regulation by IHF. Such a relation was formerly identified for the two NAPs FIS and H-NS, leading to the definition of eleven “stress-response” domains (CODOs), denoted d1 to d11, that exhibit distinct DNA physical properties, regulation by NAPs, and coherent response to growth conditions and environmental stresses [11, 13].

**Fig. 10:**
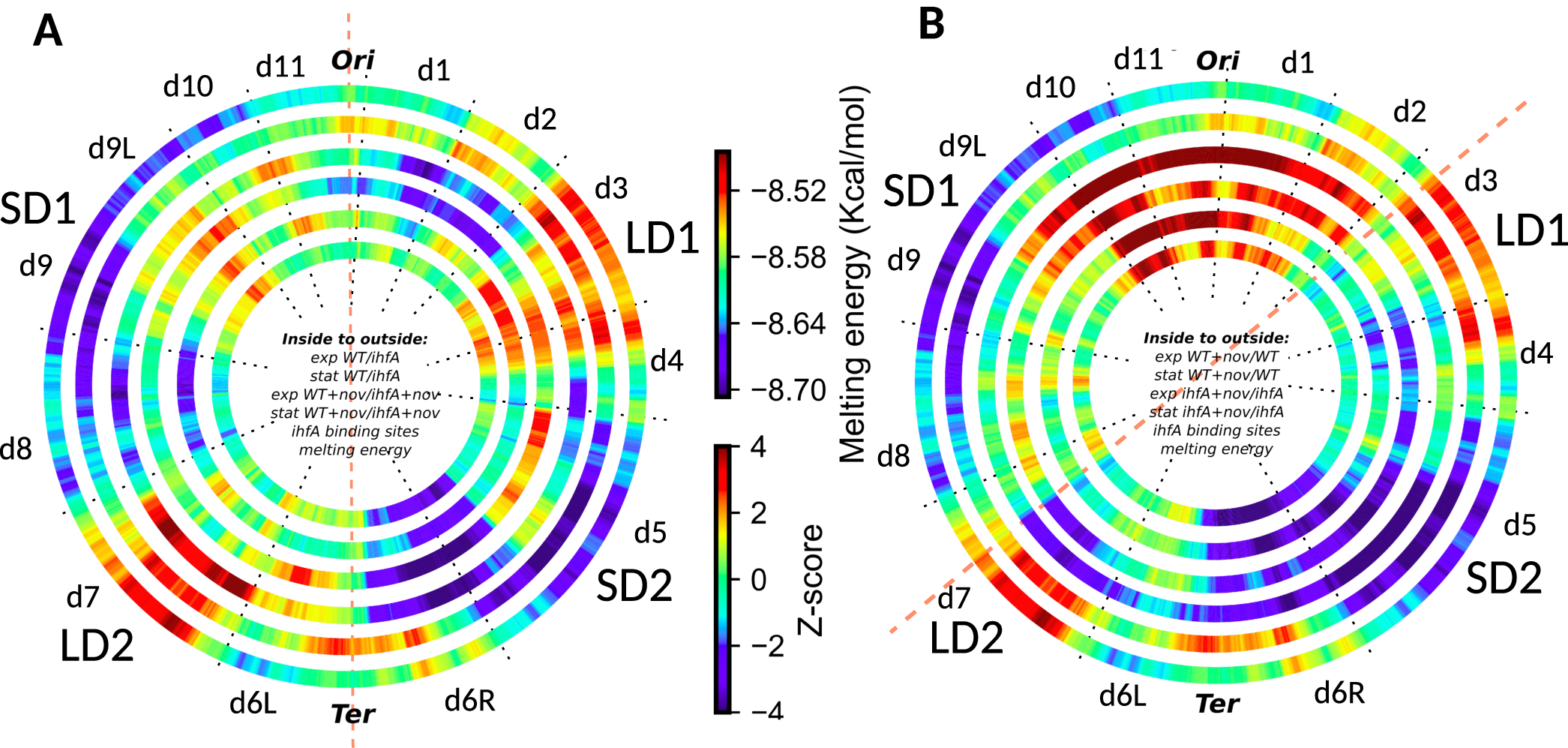
Global organisation of IHF transcriptional regulation in chromosomal domains. **(A)** Four inner rings: inter strain comparison (WT vs *ihfA*) in different growth phase (exponential or stationary phase) and supercoiling conditions (absence or presence of novobiocin). Red and blue colors indicate a significantly enriched density of activated or repressed genes, respectively. Outer ring: distribution of DNA thermodynamic stability of the *D. dadantii* 3937 genome. Second outer ring: local enrichment in predicted IHF binding sites. All quantities are computed over 500 kb windows. **(B)** Four inner rings: intra strain comparisons. Densities of novobiocin-induced (red) or -repressed (blue) genes in wild-type or *ihfA* mutant cells, in either exponential or stationary phase. Same colour code as A. Two outer rings: same as A. Note that the CODOs on both sides of OriC respond uniformly to DNA relaxation (B) while differently to IHF (A), the left side being more active in presence of IHF, especially under conditions of DNA relaxation. On both sides of OriC end, IHF counteracts the expansion of the DNA relaxation-induced activation into d2 and the OriC proximal half of d9 (d9L) in a growth phase-independent manner. This activation of d2 in *ihfA* mutant is fully consistent with the repression of this CODO under conditions of DNA relaxation when the *ihfA* mutant is compared to the wild type (compare d2, two middle rings in A and B). This domain is enriched for putative IHF binding sites, and is activated by DNA relaxation is precluded by IHF. On DNA relaxation IHF also counteracts the activation of the OriC proximal half of d9 (d9L, two middle rings in B), but no repression by IHF is observed (compare d9L, two middle rings in A). Note that at the Ter ends of both WT and *ihfA* mutant strains, DNA relaxation represses the region around d6R/d5 boundary independent of the growth phase and IHF (four inner rings in B); similarly, the adjacent d6L and d7 are repressed DNA relaxation but only in absence of IHF (two middle rings in B), leading to loss of asymmetry observed at the Ter end of wild-type cells. CODO d7 is activated by IHF under conditions of DNA relaxation in stationary phase (fourth ring from inside in A) and is repressed on DNA relaxation in *ihfA* mutant (compare fourth ring from inside in B), suggesting that this CODO is rescued from repression by DNA relaxation at high concentrations of IHF in stationary phase (Ali Azam et al. 1999). CODO d5 is activated by IHF on DNA relaxation (third ring from inside in A) and repressed by DNA relaxation in the absence of IHF during exponential growth (third ring from inside in B). Thus, also d5 appears to be rescued from inactivation at low DNA superhelical densities by IHF at a specific growth stage. Note however, that in contrast to d7 enriched for putative IHF binding sites, d5 is largely overlapping with the thermodynamically stable region SD2, poor in IHF binding sites.

The four inner rings show the genomic distributions of significantly up-regulated (red) and down-regulated (blue) genes, either between WT and *ihfA* in various conditions (inter-strain comparison, A), or during a novobiocin shock (intra-strain comparison, B). The inter-strain comparison (Fig. 10A) clearly shows that the regulatory action of IHF is not homogeneous, but organised into extended regions that largely coincide with the previously identified CODOs. This regulation by IHF is often independent of growth phase and DNA supercoiling, such as the activation of d9L and a large part of d3 extending over the d3/d4 border, as well as the repression of d6R extending over the d5/d6 border, all these effects being possibly related to the distinct thermodynamic properties of the affected regions. Interestingly, the Ter region (CODO d6) was split in two parts, showing a growth phase-independent difference in the expression level of its left (d6L) and right (d6R) halves (Fig. 10A, red dashed line). Some regions are regulated specifically in stationary phase, but independently of the DNA supercoiling level (d8, d10/d11 border), possibly reflecting a specific binding of IHF at the higher cellular concentration reached in that phase. Conversely, d2 is specifically repressed by IHF under conditions of DNA relaxation, independent of the growth phase. Finally, under conditions of DNA relaxation d7 is activated only in stationary phase, whereas d5 is activated only in exponential phase.

The intra-strain profile (Fig. 10B) of wild type cells was fully consistent with the previously observed pattern [11] and demonstrated that relaxation of DNA regulates the transcriptome in a remarkably symmetrical way, activating the chromosomal OriC end (d10, d11, d1 and the OriC-proximal half of d2) and repressing the Ter end (the region around the d6/d5 boundary) independently from the growth phase (two innermost rings). Interestingly, in the *ihfA* mutant, although the global pattern was similar to that of the WT, the regulation patches extended much further away from the Ori/Ter axis abutting in the right and left arms at the two AT-rich regions LD1 and LD2 respectively, where the IHF regulation appears strongly correlated to the thermodynamic properties of the genomic sequence.

The comparative analysis of the inter-and intra-strain transcript patterns has a potential to reveal the combinatorial effects of NAPs and DNA supercoiling [25]. Overall, we observed a conspicuous asymmetry in the activity of the left and the right chromosomal arms due to IHF-dependent opposite effects on transcription, especially under conditions of DNA relaxation (compare the two innermost and the two middle rings in Fig. 10A). In contrast, under conditions of DNA relaxation in *ihfA* mutant (intra-strain pattern, Fig. 10B) the extension of the genomic transcription in both directions from the OriC and Ter ends makes the genomic expression pattern more uniform (Fig. 10B, the two middle rings). The genome appears divided into two, predominantly activated and predominantly repressed halves, delimited by the thermodynamically labile LD1 and LD2 regions strongly enriched for putative IHF binding sites (dashed red line in Fig. 10B). Thus, it appears that IHF is required to delimit the boundaries precluding the spatial expansion of the supercoiling response in the genome (for additional details see the legend to Fig. 10). The expansion of the spatial transcription pattern from OriC/Ter in both directions observed in *ihfA* mutant on DNA relaxation might be due to cancellation of the IHF suppression of lagging strand usage.

## Discussion

The aim of this study was to determine the effect of the nucleoid-associated protein IHF on pathogenic growth and global gene expression in the model of *D. dadantii*. IHF is a component of an overarching network comprising DNA topoisomerases and the NAPs, a small class of highly abundant proteins serving both as chromatin shaping factors and as global regulators of genomic transcription [65]. Our previous studies demonstrated that inactivation of two other representatives of this class of global regulators, FIS and H-NS, strongly attenuate the pathogenic potential of *D. dadantii* [4, 14, 66]. In this work we show that mutation of *ihfA* leading to the lack of IHF heterodimer in the cells dramatically alters the transcript profile, retards cellular growth, modulates the transcriptional response to DNA relaxation by novobiocin, impairs the expression of virulence genes and as a result, abrogates the virulence of *D. dadantii* 3937. Thus, it appears that all these NAPs acting both as sensors of environmental conditions and global transcriptional regulators [65] are directly involved in coordinating the *D. dadantii* pathogenicity function with environmental conditions. The dramatic effect of *ihfA* mutation on pathogenic potential results primarily from the inability of *ihfA* mutant to utilize pectin, an important carbon source provided by the plant host, due to impaired production and/or secretion of the plant cell wall degrading enzymes, especially the Pels. This inability is reflected in the huge difference in the amount of differentially expressed genes between the WT and *ihfA* mutant strains grown in the presence of PGA, which is degraded by wild-type cells but not by the mutant (Suppl. Table S1).

### Correspondence between phenotypes and relevant genes

The phenotypic traits affected by *ihfA* mutation in *D. dadantii* are by large similar to those reported in a recent study of *Dickeya zeae* strain lacking IHF [34]. However, there are also interesting differences. For example, the *D. zeae* cells lacking IHF demonstrate decreased motility and biofilm forming capacity. In *D. dadantii* the motility is also impaired (Fig. 2), but the formation of adherence structures is increased (Fig. 3). This latter phenotype is consistent with increased expression of genes involved in production of cellulose and exopolysaccharide (Table S2). Motility and chemotaxis are essential for *D. dadantii* when searching for favourable sites to enter into the plant apoplast, as mutants with affected flagella or chemotaxis transduction system are avirulent [67]. Flagellar and chemotaxis genes are significantly affected in the *ihfA* mutant under conditions of DNA relaxation (Table S4). This includes the *fliZ* gene (Table S2) encoding the regulator of flagellar genes involved in decision between the alternative lifestyles, namely, between flagellum-based motility and biofilm formation [68] and might explain the altered motility and formation of adherence structures observed in *D. dadantii ihfA* mutant (Fig. 2 and Fig. 3). The *pil* genes encoding the type IV pilus are down-regulated, in keeping with the impaired twitching motility of the *ihfA* mutant (Table S2 and Fig. 2). The importance of type IV pilus responsible for twitching motility in *Dickeya* pathogenicity has not been studied yet. However, type IV piliation was shown to contribute to virulence in other plant pathogens, mainly in vascular pathogens, such as *Ralstonia* and *Xylella*, where they were proposed to contribute to bacterial colonization and spread in the xylem through cell attachment and twitching motility [69].

In general, the observed phenotypic changes are closely reflected in the *D. dadantii* transcriptome. The global decrease in pectinase, cellulase and protease production in the *ihfA* mutant (Fig. 2) is consistent with the impaired expression of both their cognate secretion systems (T1SS PrtEFD for proteases and T2SS OutCDEFGHIJKLMNO for pectinases and cellulase) and their coding genes (*pelA, pelC, pelZ, pelI, pelN, celZ, prtA, prtB, prtC, prtG*) (Table S2). It is noteworthy that expression of several *pel* genes (*pelB, pelD, pelE, pelW, pelX)* is not significantly affected in non-inducing conditions (Table S2) but expression of these genes is strongly decreased in the presence of inducer PGA (Table S1). The *pel* genes are regulated by a large number of global and dedicated transcription factors including H-NS, FIS, CRP, PecT, PecS, KdgR, MfbR and by the RsmA/RsmB post-transcriptional regulatory system [2]. This requirement of complex regulation likely reflects the adaptation of the bacterial lifestyle to adverse conditions of growth in various hosts, demanding a fast and reliable production of virulence factors. Expression of several regulatory genes including *fis, pecT, mfbR*, as well as the *rsmB* gene encoding regulatory sRNA, is controlled by IHF (Table S2). The dedicated transcriptional repressor PecT is repressed by IHF, and its overproduction might contribute to the plant cell wall-degrading enzyme defective phenotype of *ihfA* mutant. Indeed, a similar phenotype linked to PecT de-repression was observed in *hns* mutant [70]. The decreased expression of *fis* and *mfbR* in the *ihfA* background (Table S2) might also affect the nucleoprotein complex formed at the *pel* promoters so that it is incapable to sustain *pel* expression [71]. Indeed, MfbR is known to activate genes encoding plant cell wall-degrading enzymes in response to alkalinisation of the apoplast during the advanced stage of infection [72]. Similarly, the down-regulation of *rsmB* gene observed in *ihfA* mutant can also explain the decreased plant cell wall-degrading enzyme production (Table S2). Indeed, RsmB is a small RNA involved in the post-transcriptional control of the RNA-binding protein RsmA, which turns down the production of pectate lyases by binding directly to the *pel* mRNAs [73]. RsmB carries multiple RsmA binding sites and, therefore, titrates RsmA away from its mRNA targets [74].

The global decrease in siderophore production observed in the *ihfA* mutant (Fig. 3) is correlated with the down-regulation of the achromobactin biosynthesis operon (*acsF-acr-acsDE-yhcA-acsCBA*) as well as the *cbrABCD* operon encoding the ABC permease for ferric achromobactin (Table S2). At the same time, the *fct-cbsCEBAP* and *cbsH-ybdZ-cbsF* operons responsible for biosynthesis of the second siderophore, chrysobactin, are up-regulated in the *ihfA* mutant (Table S2). These results suggest that achromobactin is probably better detected than chrysobactin under the growth conditions on CAS agar plate (Fig. 3). Different transcriptional signatures for these two siderophores were already observed in response to various stress conditions [3]. This may explain the rationale of producing two siderophores that may be required at different stages of infection.

Other genes related to virulence are affected in the *ihfA* mutant and contribute to its reduced pathogenic potential. Notably, expression of the *hrp* genes encoding type III secretion system and its effectors DspE, HrpN and HrpW, which suppress plant immunity and promote pathogenesis, are down-regulated (Table S2). In agreement with these findings, it was shown that IHF is required for RpoN-dependent expression of *hrpL* gene encoding HrpL, the sigma factor coordinating the expression of the *hrp* genes in *Pseudomonas syringae* and *Erwinia amylovora* [33] [75].

### IHF switches the orientation preferences of transcribed genes

We observed that lack of IHF alters the preference for spatial orientation of the transcribed genes, especially under conditions of DNA relaxation. While this local effect of IHF requires further investigation, we note that it constitutes a novel and original mechanism in DNA supercoiling-dependent regulation of transcription. Analyses of previously published microarray data [11] show that H-NS does not favour any orientation, whereas FIS only slightly favours convergent genes in a relaxed chromosome (Fig. S2 in Supplementary Materials). This moderate effect of FIS might be related to the difference in the extent of bending (∼45° for FIS and ∼180° for IHF) induced on binding the DNA. However, FIS constrains toroidal coils activating promoters that require high negative superhelicity [76] whereas binding of IHF constrains little, if any negative superhelicity [15, 77] suggesting that IHF preferentially binds relaxed or slightly positively supercoiled DNA loops. This preference for relaxed templates is consistent with two other observations. First, it was observed that IHF preferentially binds at the 3’ ends of transcription units [24] which accumulate positive superhelicity due to transcription-coupled supercoil diffusion [78] [46]. Second, while IHF, alike the structurally related NAP HU, untwists the DNA by proline intercalation, the net untwisting is significantly less for IHF [77, 79]. Furthermore, since divergent transcription is expected to increase negative superhelicity in between the genes, whereas the opposite is true for convergently organised transcription units [78] [46], the former might be especially sensitive to DNA relaxation thus favouring IHF binding under this regimen. Since IHF is known to bind DNA using both direct and indirect readout [64] and stabilise various 3D structures [22], modulation of its binding by changing supercoil dynamics resulting from adjacent transcription units may lead to profound changes in the organisation of nucleoprotein complexes and depending on torsional stress, distinctly favour writhe deformations at the expense of twist. These latter can in turn favour or inhibit the formation of twist-induced DNA denaturation bubbles required for transcription initiation, as well as other competing structural transitions [63] and thereby affect convergent and divergent transcription units in different manners. The observed modification of leading/lagging strand utilisation further suggests that this interplay between IHF and DNA supercoils may not be limited to those generated by and affecting transcription, but also related to the replication machinery.

### Lack of IHF impairs the response of CODOs

Our transcriptome analyses suggest a massive reorganization of the genomic expression patterns induced by *ihfA* mutation in *D. dadantii* cells, especially under conditions of DNA relaxation induced by novobiocin addition. DNA relaxation in *ihfA* mutant background makes the genomic expression pattern more uniform compared to the wild-type, highlighting an asymmetry in relative activities of the chromosomal Ori and Ter ends. On the other hand, the comparison of the wild-type and *ihfA* mutant cells shows a difference between the chromosomal arms which again, is augmented under conditions of DNA relaxation. This differential effect of IHF on chromosomal arms is fully consistent with the observed spatial organization of transcriptional regulatory networks along the replichores in *E. coli* [80, 81]. Thus, the regulatory impacts of IHF and DNA supercoiling appear organized along the orthogonal axes (respectively along the OriC – Ter axis, and the lateral axis) of the chromosome. The combined effect of DNA supercoiling and IHF produces a variable pattern of transcriptional activity depending on the genomic organization of thermodynamically variable sequences (SD1, LD1, SD2 and LD2) and manifested as CODOs.

Our previous work demonstrated that the major virulence and adaptation genes embedded in CODOs demonstrate a regular pattern of expression that is by and large, congruent with that of the CODOs [12], supporting the notion that the flexible organisation of CODOs harboring distinct functional groups of genes and responding to different combinations of impacts, provides a *bona fide* mechanism underlying the coordinated genetic response of the *D. dadantii* cells to environmental stress [11]. This coordinated response appears impaired in *ihfA* mutant cells. For example, the CODO d7 encodes the motility and chemotaxis functions required for the colonization of the apoplast. In the absence of IHF, these functions are repressed by novobiocin treatment independent of the growth phase and accordingly, relaxation of DNA in *ihfA* mutant down-regulates CODO d7 independent of growth phase (Fig. 10B). However, d7 is activated by IHF on DNA relaxation specifically in stationary phase (Fig. 10A, fourth ring from inside). Furthermore, the CODO d6L harbours several virulence traits including type I and III secretion systems and plant wall-degrading enzymes (proteases and xylanase). This CODO is up-regulated, albeit to different extent depending on the growth phase, in wild-type cells (i.e. activated by IHF) on DNA relaxation (Fig. 10A, third and fourth rings from inside), whereas d6L is repressed by DNA relaxation in *ihfA* mutant (Fig. 10B, compare the first and second rings to the third and fourth rings from inside), consistent with the impaired pathogenic potential of the mutant strain (Fig. 4). Notably, we observed that IHF regulates all the secretion systems underpinning the bacterial pathogenic growth, as well as the genes involved in flagellar assembly and numerous membrane-anchored proteins. During the process of transcription and co-translational export, the loops of chromosomal DNA encoding the inner membrane and/or secreted proteins become transiently anchored to the plasma membrane, providing expansion forces that affect the nucleoid structure [82]. We assume that down-regulation of the corresponding genes in *ihfA* mutant may be concomitant to a corresponding profound modification of the nucleoid configuration.

Taken together, the transcriptome pattern analysis not only allowed us to identify gross genome-wide differences between the responses of the wild-type and *ihfA* cells consistent with the global effect on virulence function, but also distinguish individual CODOs, which respond to IHF, DNA supercoiling and combinations thereof, in various ways. We propose that IHF acts as a multi-scale architectural protein with a multifaceted regulatory effect. First, at the kilobase scale of topological domains, it interacts with transcription by redistributing DNA supercoiling in a strongly gene orientation-dependent manner. Second, at the megabase-scale of macrodomains, it acts as a “domainin” protein defining the boundaries for the expansion of the transcriptional supercoiling response and thus, determining the constellations of CODOs harbouring relevant genes required for faithful response of this pathogen to environmentally induced alterations of DNA topology. Third, IHF globally affects the preference for leading versus lagging strand utilization and thus determines the default setting for the directionality of transcription in the genome, as a basis for dynamic organization of CODOs. The multifaceted effect of IHF discovered in this study indicates that integrated investigation of various coordinated regulatory impacts of the NAPs and DNA supercoiling is a promising avenue for further research.

## Data availability

RNA-Seq data have been deposited at ArrayExpress repositories E-MTAB-7650 (WT strain) and E-MTAB-9025 (*ihfA* strain).

## Supplementary Data

Supplementary data for this article may be found online.

### Supplementary Tables

For Tables S1, S2, S3, S4, genes are organized by functional categories. Each gene was designated by only one functional category and genes in the same operon were classified into the same category. Column “D” indicates the operon structure in *Dickeya*.

The columns with “O”, “+” and “-” indicate whether a gene is significantly regulated by a specific condition. “O” represents no significant regulation, “+” and “-” represent the significantly up-regulated or down-regulated genes with adjusted pvalue <0.05, respectively.

**Table S1**: Transcriptional response to PGA in WT strain and *ihfA* mutant, at transition to stationary phase

**Table S2**: Transcriptional response to *ihfA* mutation

**Table S3**: Transcriptional response to novobiocin shock in WT strain and *ihfA* mutant

**Table S4**: List of *ihfA*-differentially expressed genes under DNA relaxation only

**Table S5**: List of predicted IHF binding sites at gene promoters.

### Supplementary Fig.s

Fig. S1: Density of predicted IHF binding sites **(A)** and A/T bases **(B)** in intergenic regions between convergent, divergent and tandem genes. Error bars represent 95% confidence intervals.

Fig. S2: Transcriptional effect of *fis* **(A)** and *hns* **(B)** mutation, in absence or presence of novobiocin, depending on gene orientation. In contrast to the strong and systematic orientational effect of *ihfA*, only *fis* presents a limited comparable effect in a relaxed chromosome only. Error bars represent 95% confidence intervals.

## Acknowlegments

The authors thank Corentin Schotte, Florelle Deboudard, Elodie Kenck, Camille Villard for technical support and Hubert Charles for helpful discussions.

## Funding

This work was supported by the INSA grant to GM as visiting researcher. R.F was funded by a research allocation from the French Ministry. This work also benefited from INSA Lyon grants [BQR 2016 to S.M.]; IXXI; Agence Nationale de la Recherche [ANR-18-CE45-0006-01 to S.M.], Breakthrough Phytobiome IDEX LYON project, Université de Lyon Programme d’investissements d’Avenir [ANR16-IDEX-0005]; Centre National de la Recherche Scientifique [to S.R., F.H. W.N. and S.M.]; Université de Lyon [to S.R., F.H. W.N. and S.M.].

## Notes

### Competing Interest Statement

The authors have declared no competing interest.

