## Supplementary figures and images for "The nucleoid-associated protein IHF acts as a “domainin” protein coordinating the bacterial virulence traits with global transcription"

### figS1.pdf

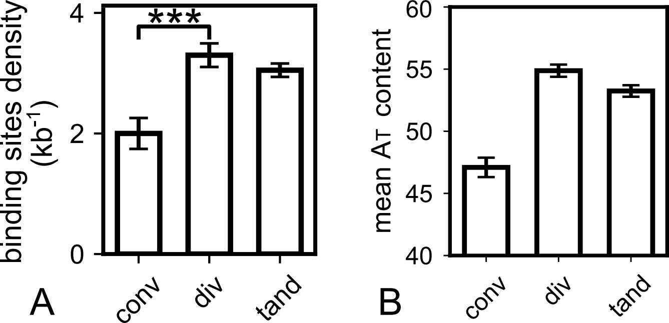

### figS2.pdf

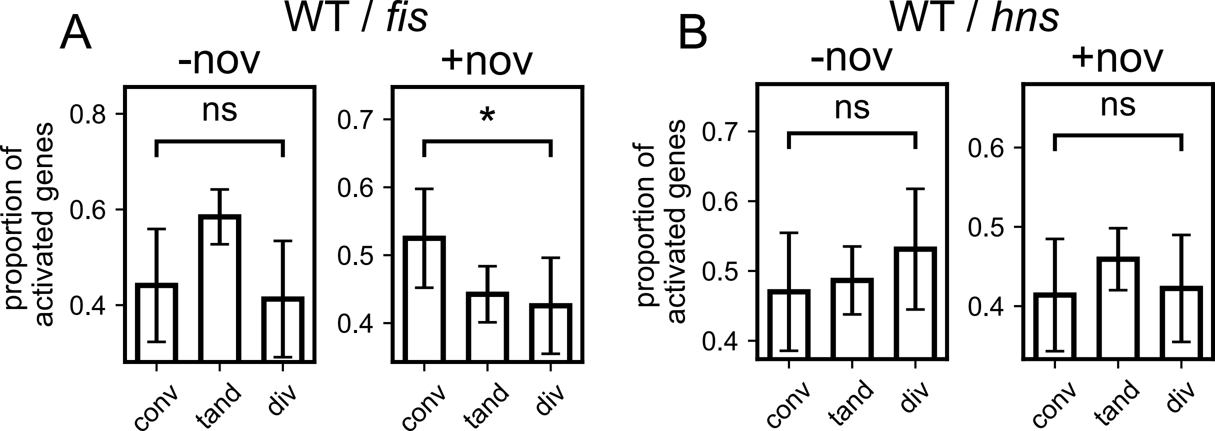
